# Entomopathogenic nematodes increase predation success by inducing specific cadaver volatiles that attract healthy herbivores

**DOI:** 10.1101/442483

**Authors:** Xi Zhang, Ricardo A. R. Machado, Cong Van Doan, Carla C. M. Arce, Lingfei Hu, Christelle A. M. Robert

## Abstract

Herbivore natural enemies, including predators, parasitoids and parasites, protect plants by regulating herbivore populations. Some parasites can increase their transmission efficiency by manipulating host behavior. Whether natural enemies can manipulate herbivore behavior to increase top-down control, however, is unknown. Here, we investigate if and how the entomopathogenic nematode *Heterorhabditis bacteriophora,* an important biocontrol agent, modulates the behavior of the western corn rootworm, *Diabrotica virgifera virgifera,* a major maize pest, and how these behavioral changes affect the capacity of the nematode to control the rootworm. We found that healthy rootworm larvae are attracted to nematode-infected cadavers shortly before the emergence of the next generation of nematodes. Nematode-infected rootworms release distinct volatile bouquets, including butylated hydroxytoluene (BHT), which attracts rootworms to infected cadavers. In a soil setting, BHT attracts rootworms and reduces nematode resistance, resulting in increased infection rates and rootworm mortality as well as increased nematode reproductive success. Five out of seven tested insect species were found to be attracted to nematode-infected conspecifics, suggesting that attraction of healthy hosts to nematode-infected cadavers is widespread. This study reveals a new facet of the biology of entomopathogenic nematodes that increases their capacity to control a major root pest by increasing the probability of host encounters.

## INTRODUCTION

Herbivore natural enemies such as predators, parasites and parasitoids play a key role in terrestrial ecosystems by reducing herbivore abundance (1). Biological control relies on this form of top-down control to protect crops from herbivores (2). In order to exert their effects, herbivore natural enemies need to make contact with their hosts. Natural enemies have evolved various behavioral strategies to maximize their chance to encounter herbivores (3–6). Predators and parasitoids for instance can use herbivore-induced plant volatiles to locate herbivores (7). Herbivores on the other hand can detect and actively avoid contact with natural enemies (8). The interplay between behavioral adaptations of herbivores and natural enemies is likely to be an important determinant for the success of herbivore natural enemies and their capacity to suppress herbivore pests.

A key step in the life of many herbivore natural enemies is the acquisition of new hosts once the old host is exploited. Predators and parasitoids acquire new hosts by hunting, ambushing and trapping them. Parasites with indirect life cycles can also facilitate the transfer to new hosts through host manipulation strategies, including changes in color, smell and behavior of their current hosts to attract alternate hosts (9–12). How parasites with direct life cycles (i.e. involving a single host) can facilitate transmission from exploited hosts to new healthy hosts is less well established (13). Recent studies show that insect bacterial pathogens can induce to changes in volatile emissions in infected individuals, which results in the attraction of non-infected individuals (14). Whether predators, parasitoids and multicellular parasites with direct life cycles can use volatiles to attract additional hosts or prey remains to be determined. Furthermore, whether behavioral manipulation of herbivores by natural enemies can enhance top-down control of herbivores is unclear.

Entomopathogenic nematodes (EPNs) are important biological control agents of insect herbivores (15, 16). EPNs can locate their herbivores using volatile cues that are emitted by herbivores and herbivore-infested plants (17). Once an EPN comes into contact with a compatible host, it penetrates it and injects entomopathogenic symbiotic bacteria, which kill the insect (18). The EPN then feeds on bacteria and infected host-tissues and multiplies within the cadaver. Eventually, a new generation of infective juveniles emerges from the cadaver and begins searching for new hosts. A crucial factor that determines the success of EPNs is their transmission efficiency from exploited to healthy hosts (19). As EPNs are much less mobile than their hosts, they may have evolved host manipulation strategies to increase the probability of host encounters. Conversely, herbivores may effectively avoid entomopathogenic nematodes by detecting their presence and avoiding them. So far, the impact of EPNs on host behavior has not been studied in detail.

Here, we investigated how infection by the EPN *Heterorhabditis bacteriophora* influences the behavior of healthy herbivores. We first studied the behavior of the western corn rootworm (WCR, *Diabrotica virgifera virgifera),* a major root pest of maize who occurs sympatrically with *H. bacteriophora* and is the target of EPN-based biological control programs. WCR can use plant toxins to repel *H. bacteriophora* (20), but successful biological control of WCR through *H. bacteriophora* has nevertheless been reported (21, 22). Through a series of behavioral experiments, we demonstrate that healthy WCR larvae are attracted to EPN-infected cadavers. We identify a volatile that is specifically released from infected cadavers and attracts WCR larvae. We use this volatile to assess how the attraction of healthy hosts affects the capacity of EPNs to infect and kill WCR in the soil. Finally, we determine the impact of EPN infection on the volatile release and attraction of different insect species to test whether this phenomenon may be widespread. Collectively, these experiments reveal a novel facet of EPN biology that enhances their capacity to infect and kill insect herbivores.

## RESULTS

### Western corn rootworm larvae are attracted to nematode-infected cadavers

To explore how WCR responds to the presence of nematode-infested conspecifics, we infested root-feeding WCR larvae with *H. bacteriophora* entomopathogenic nematodes (EPNs). We then measured the attractiveness of the maize+WCR+EPN complexes at different time points over 96 hr using belowground olfactometers. Maize+WCR+EPN complexes were attractive to WCR 48 hr after WCR and EPN application (Fig. 1A). The attraction coincided with high root consumption by WCR (Fig. 1B) and high production of the WCR attractant (*E*)-β-caryophyllene (23, 24) by the attacked maize roots (Fig. 1C). At this time point, approx. 30% of the WCR larvae were infected with EPNs (Fig. 1D). The attractive effect of maize-WCR-EPN complexes disappeared at 72 hr but reappeared at 96 hr (Fig. 1A). The increased recruitment of WCR 96 hr post infestation was unexpected, as at this time point 95% of WCR larvae were infected and killed by EPNs (Fig. 1D), and root removal and root (*E*)-β-caryophyllene emissions had decreased markedly (Fig. 1B and 1C).

**Figure 1.**
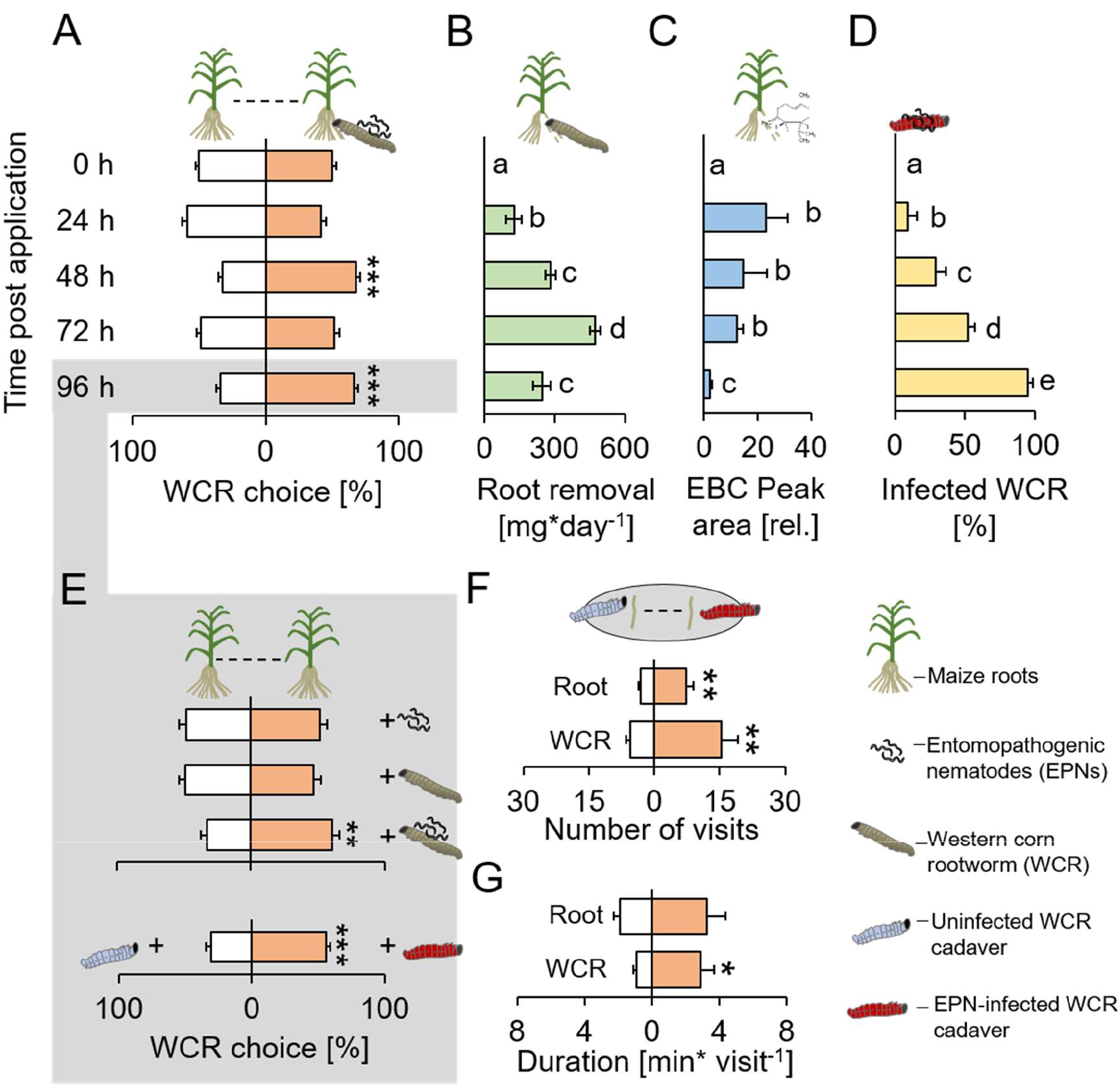
Root herbivore recruitment dynamics of plant-herbivore-natural enemy complexes reveal that herbivore cadavers infected by entomopathogenic nematodes attract healthy herbivores. **A.** Proportions (mean ± SEM) of western corn rootworm (WCR) choosing between healthy maize plants and maize plants infected with conspecifics and entomopathogenic nematodes (EPNs) in belowground olfactometers. WCR choice was measured 0 hr, 24 hr, 48 hr and 96 hr after infection (n=45). **B.** Root removal (mean ± SEM) by WCR larvae 0 hr, 24 hr, 48 hr and 96 hr after infection (n=5-8), **C.** (*E*)-β-caryophyllene (EBC) production (mean ± SEM) of maize roots 0 hr, 24 hr, 48 hr and 96 hr after infection (n=3-5), **D.** WCR infection by EPNs (mean ± SEM) 0 hr, 24 hr, 48 hr and 96 hr after infection (n=8). **E.** Proportions (mean ± SEM) of WCR larvae choosing healthy plants or plant+WCR+EPN complexes (n=20), healthy plants or WCR-infested plants (n=20), healthy plants or plant+WCR+EPN complexes (n=20), caged uninfected or EPN-infected WCR cadaver (n=33). Larval preference was assessed in belowground olfactometers 96 hr post infection. **F-G.** Number and duration of visits (mean ± SEM) of WCR larvae exposed to uninfected and EPN- infected WCR cadavers in the presence of maize root pieces (n=6). Stars indicate significant differences (*: p<0.05, **: p<0.01, ***: p<0.001).

These experiments show that the interaction between maize, WCR and EPNs results in dynamic changes in WCR recruitment over time, with maize+WCR+EPN complexes becoming attractive as WCR infection by EPNs progresses.

To better understand the factors that render plant-herbivore-nematode complexes attractive to WCR 96 hr post infection, we quantified WCR recruitment to WCR and EPNs individually and in combination. Plants in the presence of WCR or EPNs alone were not attractive to WCR than plants alone at this time point. However, plants in the presence of WCR under EPN attack were attractive to WCR larvae (Fig. 1E). We next tested whether the cadavers of EPN-infected WCR larvae attract WCR by putting infected cadavers and uninfected WCR larvae into small filter paper cages and burying them beneath individual maize plants. WCR larvae preferred EPN-infected WCR cadavers over uninfected cadavers (Fig. 1E). Time course analysis revealed that the attraction to EPN- infected cadavers was strongest 96 hr after infection, shortly before the emergence of infective juveniles (Fig. S1). To better understand how WCR larvae respond to the presence of EPN-infected cadavers, we performed additional behavioral experiments in petri dishes (25) using EPN-infected and uninfected WCR cadavers and small root pieces to provide plant background odors. WCR larvae visited infected cadavers more often than uninfected cadavers (Fig. 1F) and spent more time per visit on infected cadavers (Fig. 1G). The number of root visits was also increased for roots that were close to infected cadavers (Fig. 1F). WCR larvae did not show any preference for uninfected or EPN-infected cadavers in the absence of plant roots (Fig. S2A). We hypothesized that this may either be due to plant background odors which are required to elicit WCR search behavior, or due to plant-mediated attraction, where EPN-infected WCR cadavers render roots more attractive to WCR. To test the second hypothesis, we exposed maize roots to uninfected and EPN-infected cadavers, removed the cadavers and evaluated WCR choice. WCR larvae did not show any preference for the different roots (Fig. S2B). Together, these experiments demonstrate that EPN infection of WCR directly increases volatile-mediated recruitment and cadaver contact of healthy foraging WCR larvae.

### Nematode-infection induces volatile release from herbivore cadavers

To identify possible volatile cues that may attract WCR to EPN-infected cadavers, we performed headspace analyses of infected an uninfected WCR cadavers. GC-FID analysis revealed no significant difference in CO_2_ emissions between uninfected and EPN-infected WCR cadavers (Fig. S3).

Headspace solid phase micro extraction (SPME) and GC-MS followed by automated alignment and peak picking revealed 279 distinct volatile features, including 15 features that were exclusively detected in the headspace of infected cadavers (Fig. 2A). Principal component analysis (PCA) revealed a clear separation of volatile profiles from infected and uninfected cadavers along PC axis 1 (Fig. 2B). A single volatile eluding at 18.23 min explained 45.6 % of the variability of axis 1 and was exclusively present in the headspace of infected cadavers (Fig. 2C). Additional manual integration and relative quantification of the 11 highest peaks in the headspace chromatograms revealed that 9 out of 11 integrated peaks were emitted in higher quantities by infected cadavers (Fig. 2D). The highest peak, at retention time 18.23 min, was exclusively present in chromatograms of infected cadavers. Based on comparisons of mass spectra and co-injection of a synthetic standard, the compound was identified as butylated hydroxytoluene (BHT). A single infected WCR cadaver was found to release up to 5 ng of BHT per hr (Fig. S4A). Emission kinetics showed that BHT starts being released 72 hr after EPN infection (Fig. S4A), which corresponds to the onset of increased WCR recruitment to EPN-infected cadavers (Fig. S1). Therefore, further investigations focused on BHT as a potential volatile attractant of WCR. BHT has originally been described as a synthetic antioxidant (26), but is also naturally produced by cyanobacteria, algae, and fungal pathogens (26, 27). To test whether the EPN endosymbiontic bacterium *Photorhabdus laumondii* subsp. *laumondii* (28) may be responsible for BHT production, we injected it into WCR larvae directly, which resulted in visual infection symptoms and mortality similar to EPN infection. No BHT release from *P. laumondii* infected WCR cadavers was detected (Fig. S4B). *P. laumondii* grown *in vitro* did not release any BHT either (Fig. S4B). We also did not detect any BHT release from EPNs alone (Fig. S4B) or uninfected WCR cadavers (Fig. 2). Instead, BHT was exclusively detected in EPN-infected WCR cadavers (Fig. S4B). These results imply that BHT release is specific to infection of WCR by EPNs.

**Figure 2.**
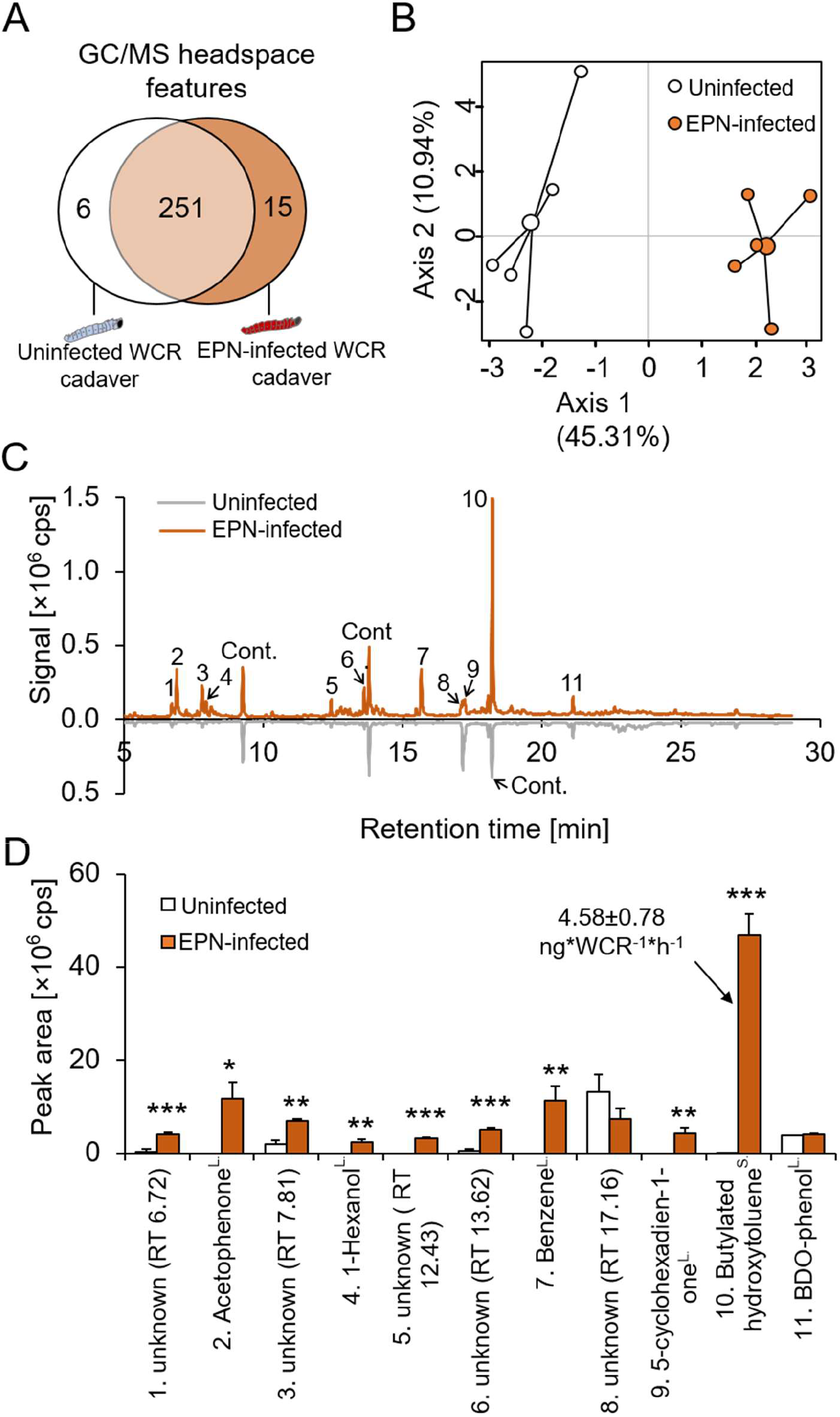
Butylated hydroxytoluene is specifically emitted by western corn rootworm larvae that are infected by entomopathogenic nematodes. **A.** Venn diagram showing the numbers of overlapping and non-overlapping GC-MS headspace features of uninfected (white) and EPN-infected (brown) western corn rootworm (WCR) cadavers (n=5). **B.** Principal component analysis (PCA) of volatile emissions of uninfected (white) and EPN-infected (brown) WCR cadavers (n=5). **C.** Representative GC-MS volatile profile of uninfected (white) and EPN-infected WCR larvae (brown). 1, 3, 5, 6, 8: unknown; 2: aceptophenone; 4: 1-hexanol; 7: benzene; 9: 5-cyclohexadien-1-one; 10: butylated hydroxytoluene (BHT); 11: 2,6-bis (1,1-dimethylethyl)-4-(1-oxopropyl) phenol; Cont.: contamination. cps: count per second. **D.** Volatile peak areas (mean ± SEM) of uninfected (white) and EPN-infected WCR larvae (brown) (n=5). cps: count per second. ^L^: identification based on libraries. ^S^: identification based on pure standards. Stars indicate significant differences (*: p<0.05, **: p<0.01, ***: p<0.001).

### Butylated hydroxytoluene emitted by nematode-infected cadavers attracts healthy hosts and renders them more susceptible to nematode attack

Based on the correlation between BHT release and WCR attraction (Fig. S1 and S4), we hypothesized that BHT may recruit WCR larvae to EPN-infected cadavers. At physiologically relevant doses, synthetic BHT was attractive to WCR and elicited responses that were comparable to infected cadavers (Fig. 3A). Furthermore, BHT complementation of uninfected WCR cadavers rendered them as attractive as infected cadavers (Fig. 3A). Thus, BHT is sufficient to attract WCR to nematode infected cadavers. BHT exposure may not only increase the recruitment of herbivore hosts but may also affect their nematode resistance. To test this hypothesis, we pre-incubated WCR larvae with BHT and then measured nematode infection rates in a no-choice setup. Pre-exposure of WCR to BHT increased infection rates by 19% (Fig. 3B). By contrast, pre-exposure of EPNs to BHT did not affect their capacity to infect WCR (Fig. 3D). Interestingly however, EPNs were attracted by EPN-infected cadavers as well as BHT similar to WCR (Fig. 3C). Thus, in addition to attracting WCR larvae, BHT renders them more susceptible to EPNs and attracts EPNs themselves.

**Figure 3.**
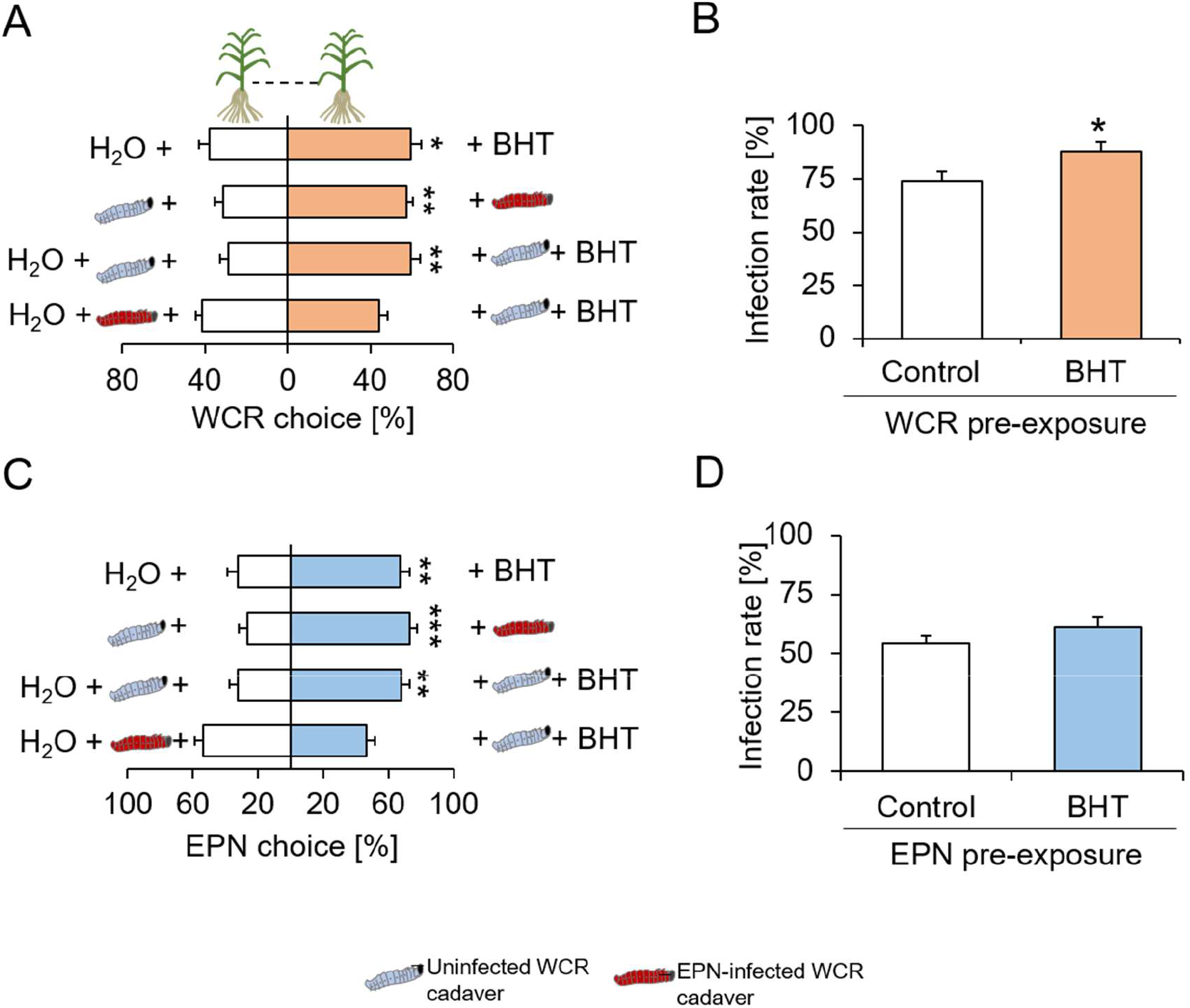
Butylated hydroxytoluene attracts herbivores and makes them more susceptible to entomopathogenic nematodes. **A**. Proportions (mean ± SEM) of western corn rootworm (WCR) larvae orienting towards BHT or H_2_O (n=20), uninfected WCR cadavers or cadavers infected by entomopathogenic nematodes (EPNs, n=10), uninfected WCR cadavers covered with BHT or H_2_O (n=15), EPN-infected WCR cadavers covered with BHT or H_2_O (n=15). **B**. WCR infection rate (Mean ± SEM) after exposure to BHT or H_2_O (n=10). **C**. Proportions (Mean ± SEM) of EPNs orienting towards *Diabrotica balteata* exudates complemented with BHT or H_2_O, uninfected WCR cadavers or cadavers infected by EPNs, uninfected WCR cadavers covered with BHT or H_2_O, EPN-infected WCR cadavers covered with BHT or H_2_O (n=20). **D**. WCR infection by EPNs (Mean ± SEM) after pre-incubation of EPNs with BHT or H_2_O for 24 hr (n=15). Stars indicate significant differences (*: p<0.05, **: p<0.01, ***: p<0.001). All treatment solutions contained 0.01% ethanol.

### Butylated hydroxytoluene increases nematode predation success in the soil

To understand how the release of BHT by EPN-infected cadavers influences EPN predation success, we conducted experiments in soil arenas. Infective juvenile EPNs were added with or without BHT to different sides of the arenas, and healthy WCR larvae were released in the middle (Fig. 4A). Significantly more WCR larvae were recovered in the vicinity of BHT presence (Fig. 4B), thus confirming the attractive effect of BHT in the soil. The proportion of EPN-infected WCR larvae was three times higher on the BHT supplemented side (Fig. 4C), and three times more nematodes of the next generation emerged from the BHT side (Fig. 4D). Thus, the increase in BHT- mediated WCR recruitment is associated with increased predation success and total offspring production of entomopathogenic nematodes.

**Figure 4.**
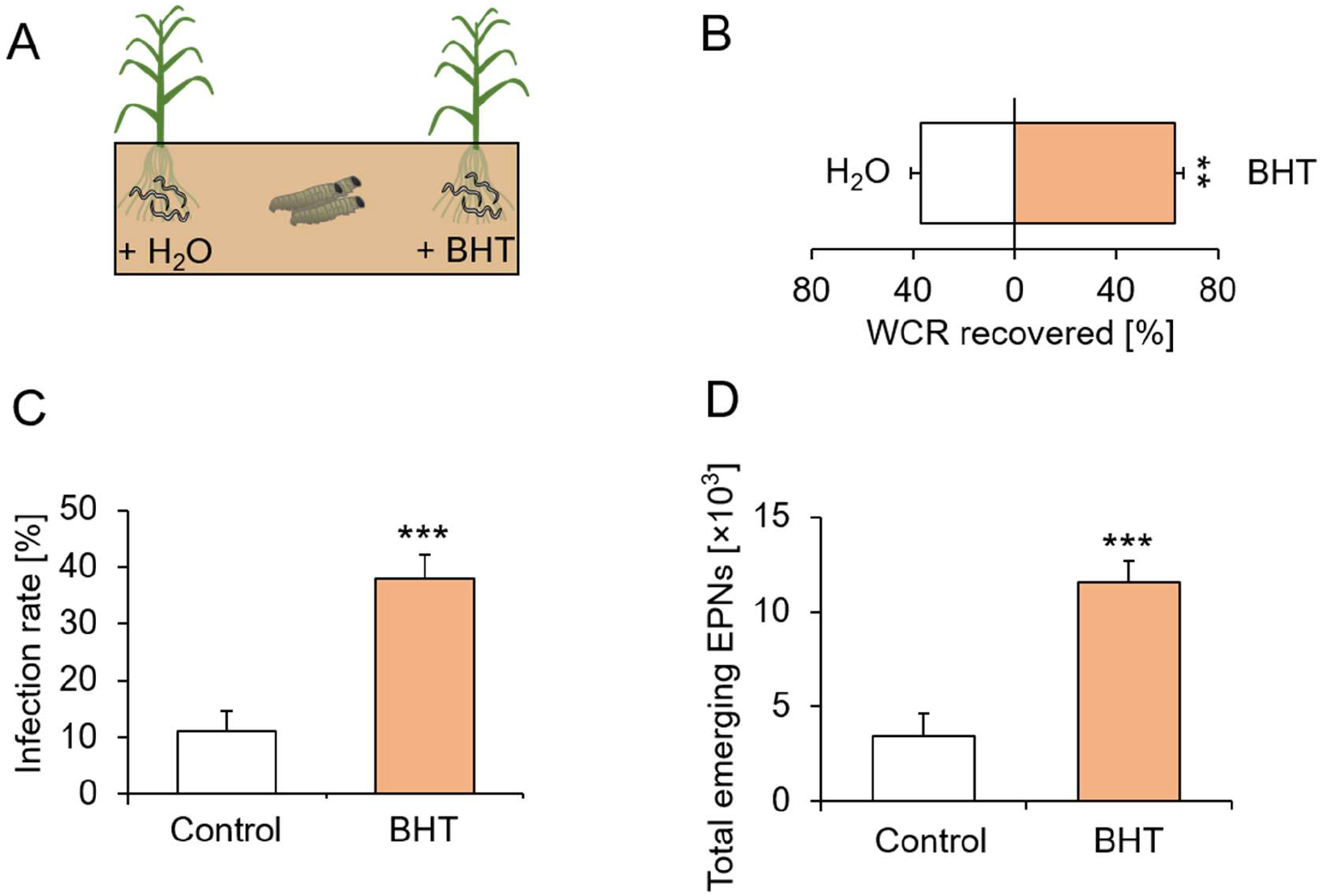
Butylated hydroxytoluene increases herbivore recruitment, predation success and fitness of entomopathogenic nematodes in the soil. A. Visual representation of experimental setup. Entomopathogenic nematodes were applied on both sides of the arenas, and each side was either watered with BHT or H_2_O. Eight western corn rootworm (WCR) larvae were then released in the middle and recollected after five days (n=12). B. Proportions (Mean ± SEM) of WCR larvae recovered from each side (n=12). C. WCR infection rates (Mean ± SEM) on each side (n=12). D. Total number of EPNs (Mean ± SEM) emerging from the WCR larvae on each side (n=12). Stars indicate significant differences (**: p<0.01, ***: p<0.001).

As EPNs themselves are also attracted by BHT (Fig. 3C), we measured whether EPNs from the control side of the arenas may have moved to the BHT side by using a *Galleria melonella* baiting approach (29). EPNs to one side of the arena led to *G. melonella* infection of the other side, and the infection rate was slightly increased when BHT was added (Fig. S5). Thus, the observed increase in WCR predation is likely to be the result of increased WCR recruitment, increased WCR susceptibility and increased EPN recruitment.

### Recruitment of healthy hosts to nematode infested cadavers is widespread and associated with the induction of species-specific volatile profiles

As *H. bacteriophora* is a generalist parasite that can infect many other insect species above and below ground, we investigated whether *H. bacteriophora* infection also increases the recruitment of healthy hosts in other insect species. Five of the seven tested species, including *Diabrotica virgifera* (WCR), *D. balteata, Tenebrio molitor, Drosophila melanogaster* and *Spodoptera littoralis* were attracted to cadavers of EPN-infected conspecifics (Fig. 5A). Only the larvae of the honeycomb moth *Galleria melonella* larvae, which are not typically exposed to EPNs in nature, preferred non-infected over infected cadavers. Thus, the attraction of healthy insect hosts to infected conspecifics is widespread among EPN host species.

**Figure 5.**
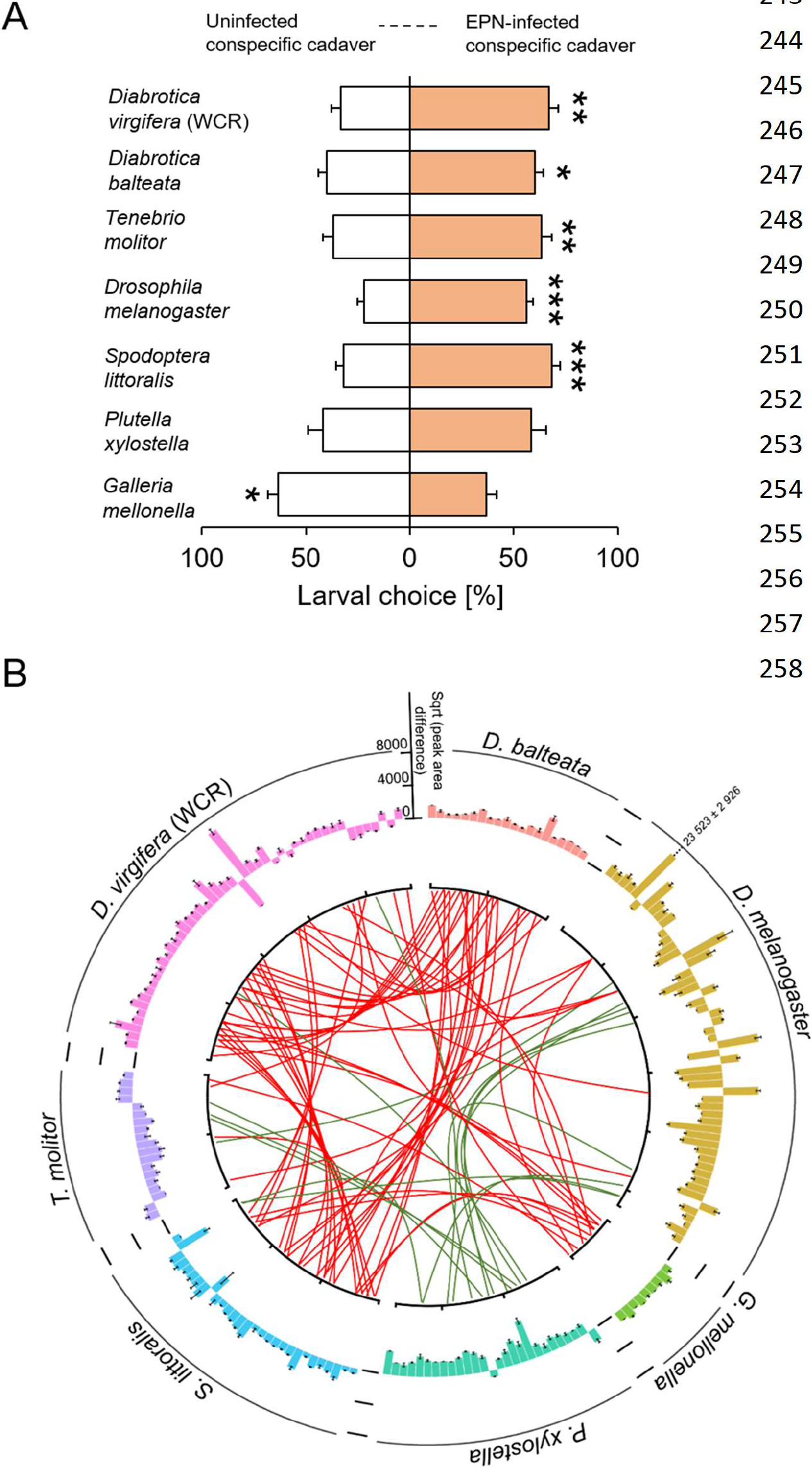
Attraction to cadavers infected by entomopathogenic nematodes is a widespread and associated with specific volatile bouquets. **A.** Proportions (mean ± SEM) of larvae orienting towards conspecific cadavers that were uninfected or infected by entomopathogenic nematodes (EPNs, n=10). A total of seven species were tested: *Diabrotica balteata* (n=19), *Tenebrio molitor* (n=16), *Drosophila melanogaster* (n=14), *Spodoptera littoralis* (n=18), *Plutella xylostella* (n=16) and *Galleria mellonella* (n=10). Stars indicate significant differences (*: p<0.05, **: p<0.01, ***: p<0.001). **B.** Differentially regulated volatiles (p<0.10) released by EPN-infected cadavers compared to uninfected cadavers of the different insect species (square root transformed data mean ± SEM, n=10-19). Red lines indicate volatiles that are emitted in higher amounts by EPN-infected cadavers than by uninfected cadavers. Green lines indicate volatiles that are emitted in lower amounts by EPN- infected cadavers than by uninfected cadavers.

To determine whether the attraction of the different insect species to infected cadavers can be explained by BHT release, we screened volatile emissions of healthy and infected cadavers of all tested insect species. EPN infection significantly altered the volatile bouquets of all insects (Fig. 5B). Surprisingly however, the induced volatiles differed substantially between species. No commonly induced volatiles were found across all species or across the six species that were attracted to infected cadavers. BHT was only released from EPN-infected WCR and *D. melanogaster* cadavers. From these experiments, we conclude that the attraction of healthy conspecifics to EPN-infected cadavers likely involves the emission of distinct, host-specific attractants.

## DISCUSSION

Natural enemies reduce herbivore populations and thereby contribute to the dominance of plants in terrestrial ecosystems and to high plant yields in agriculture. Parasites with indirect life cycles are well known to be able to increase their transmission by manipulating host behavior (30, 31), but the prevalence and importance of this phenomenon in parasites with direct life cycles, including herbivore natural enemies, as its impact on top-down control of herbivores remains largely unexplored. Here, we show that nematode-infection triggers the release of volatiles from the cadavers of herbivorous insects, resulting in the attraction of healthy herbivores, increases infection rates and increased nematode reproduction. Below, we discuss the mechanisms and implications of these findings.

Parasites have developed fascinating and diverse strategies to manipulate their hosts and thereby increase their fitness (31–33). Here, we show that entomopathogenic nematodes increase their predation success by inducing the release of volatiles from infected host cadavers. These volatiles attract healthy herbivores and reduce their capacity to resist nematode attack. Thus, when the next generation of infective juvenile nematodes emerges from the exploited cadaver, they have a higher chance of increasing healthy hosts, which boosts their chances of survival and reproduction. Other nematodes also cue in on these volatiles, which may also increase their chance of encountering additional hosts. Together, these phenomena markedly increase top-down control of herbivores in the soil, as shown here for an important agricultural pest, the western corn rootworm. Earlier work shows that entomopathogenic nematodes also follow plant volatiles (34), which serve as aggregation cues to the western corn rootworm (23, 24, 35). The capacity to attract healthy hosts represents a new facet of EPN biology that may explain why entomopathogenic nematodes can control the western corn rootworm in the field despite the fact that the insect sequesters plant toxins for self-defense (20–22).

EPN-infected WCR cadavers release a distinct bouquet of volatiles, including butylated hydroxytoluene (BHT). Butylated compounds such as BHT are uncommon in nature, and naturally produced BHT has so far only been found in a handful of microorganisms (26, 36). We found that BHT is specifically released from EPN-infected WCR cadavers, and that it is sufficient to elicit WCR behavior similar to EPN-infected cadavers. How BHT is produced in the cadavers requires further study. Digestion of the larvae by symbiotic bacteria of the nematodes is not sufficient to elicit BHT release, suggesting that nematode-specific factors are required. Entomopathogenic nematodes produce a variety of proteins to overcome and digest their insect hosts (37), and it is probable that these proteins interact with host-derived metabolites to form BHT. BHT is a radical scavenger that is used as a food additive and synthetic analog of vitamin E (38). Thus, the production of BHT may have additional benefits to the nematodes, for instance by preserving the herbivore cadavers as they are consumed. From an applied point of view, BHT may represent a cost-effective synthetic substance that could be applied as a bait that attracts the western corn rootworm and its natural enemies.

Parasites typically attract healthy hosts by hijacking adaptive behavioral responses. The flatworm *Leucochloridium paradoxum* for instance modifies the eye stalks of snails to resemble caterpillars, which prompts birds to attack the eyes, thus allowing the flatworm to be transmitted to its primary host (39). Furthermore, the bacterial pathogen *Pseudomonas entomophila* triggers the release of aggregation pheromones from infected *Drosophila melanogaster*, which attracts healthy flies and thus enhances pathogen dispersal (14). We show that entomopathogenic nematodes can use volatile such as BHT to attract healthy rootworm larvae in the soil. Why the rootworm larvae are attracted by the volatiles of infected cadavers is currently unclear. Given that approaching an infected cadaver bears a substantial mortality risk due to the presence of infective juveniles and the suppression of immunity by volatiles such as BHT, it is unlikely that this behavior is adaptive for the herbivore itself. Based on the current literature, it seems more likely that following volatiles such as BHT is beneficial for the rootworm in a different context. Because WCR larvae and adults are attracted to and use certain aromatic compounds for host selection (25, 40), one hypothesis is that BHT attracts the larvae either by interacting with the receptors of compounds involved in host location or by mimicking their activity. Such effects were for instance reported for volatile odorants blocking CO_2_ receptors and responses in fruit flies and mosquitoes (41, 42).

Even though the benefits of attracting healthy rootworms for entomopathogenic nematodes seems evident, whether this is a true form of manipulation requires further mechanistic, evolutionary and ecological insights. As *H. bacteriophora* is a generalist with a broad host range, we hypothesized that the nematode should be able to induce attractive volatiles in a wide variety of hosts in order to benefit from this trait. Indeed, nematode-infestation triggered attraction of healthy conspecifics in five out of seven tested insect species, suggesting that this phenomenon is widespread and may benefit *H. bacteriophora* in the presence of different insect host species. Surprisingly, our analyses of the volatile blends that are emitted upon infection by the different insects revealed a high degree of specificity, with each insect producing a different, attractive volatile blend, with little overlap between the different species. It is tempting to speculate that *H. bacteriophora* may have the capacity to adjust its capacity to induce attractive volatiles to the host it invades to maximize the attraction of healthy conspecifics. Different insect species are attracted by different volatiles (43–45), which makes such an approach necessary if it is to work across different hosts. A better understanding of the proximal mechanisms of volatile induction by nematode infection would help to shed light on this hypothesis (46).

In conclusion, this study demonstrates that infection with entomopathogenic nematodes triggers the release of volatiles that are attractive to healthy hosts and suppress their nematode resistance, which increases predation success and top-down control of a herbivore pest. The finding that nematode infection increases the recruitment of healthy hosts across different insect species suggests that this phenomenon is widespread and may contribute to shaping the interactions between insects and their natural enemies in nature and in the context of the biological control of soil-borne insect pests.

## METHODS

### Biological resources

Maize plants (*Zea mays* L.; variety ‘Akku’, Delley semences et plantes SA, Swizerland) were grown in a greenhouse (23 ± 2 ^°^C, 60% relative humidity, 16:8 hr L/D, and 250 μmol.m^−2^.s^−1^ light). Twelve-day-old plants were used for all the experiments. Western corn rootworm (WCR, *Diabrotica virgifera virgifera* LeConte) eggs were provided by Chad Nielson and Wade French (USDA-ARS-NCARL, Brookings, SD, USA). *Diabrotica balteata* (LeConte) and *Plutella xylostella* eggs were kindly supplied by Oliver Kindler (Syngenta Crop Protection AG, Stein, Switzerland). *Drosophila melanogaster* wild type strain were obtained from institute of cell biology, University of Bern (Bern, Switerland). *Spodoptera littoralis* eggs were provided by Ted Turlings (University of Neuchâtel, Neuchâtel, Switzerland). *Galleria melonella* and *Tenebrio molitor* larvae were obtained from a commercial ventor (Fischereibedarf Wenger, Bern, Switzerland). Entomopathogenic nematodes (EPNs) were bought from Andermatt Biocontrol (Andermatt Biocontrol, Grossdietwil, CH) and reared in *Galleria mellonella* larvae as described by McMullen *et al* (47). EPN-infected insect larvae were obtained by placing larvae in solo-cups (30 mL cups, Frontier Scientific Services, Inc., USA) containing a 0.5 cm layer of autoclaved moist sand (Selmaterra, Bigler Samen AG, Thun, CHE) and 1000 infective juveniles (IJs) EPNs in 500 μL tap water. Uninfected controls were obtained by adding 500 μL tap water to the insect larvae.

### WCR preference

#### Belowground olfactometer experiments

The impact of EPN infection on WCR behavior was investigated in a series of experiments using dual-choice olfactometers as described previously (24). Briefly, different combinations of plants, EPNs and WCR larvae were placed in L-shaped glass pots. One control plant and one treated plant were connected by one glass connector closed by two Teflon connectors containing a fine mesh preventing the larvae from accessing the plant root system but allowing the spread of volatiles through the different compartments. Five third-instar WCR larvae were added to the central connector. The first choice of the larvae was monitored. Larvae remaining in the central connector longer than 15 min were recorded as “no choice”. First, WCR larvae were given the choice between a healthy maize plant and a maize plant surrounded by five conspecifics and 2000 EPNs. WCR choice was evaluated after 0, 24, 48, 72 and 96 hr (n=45 for each time point). Second, WCR preference 96 hr post infestation was further investigated by offering larvae the choice between the following combinations: (i) plant *vs*. plant+EPNs (n=20); (ii) plant *vs*. plant+WCR (n=20); (iii) plant *vs*. plant+WCR+EPN (n=20) and (iv) five WCR larvae enclosed in a filter paper cage next to a healthy plant *vs*. five EPN-infected WCR larvae enclosed in a filter paper cage next to a healthy plant (n=33). Infestation of plants with WCR larvae was realized by adding five third instar larvae to the bottom opening of the glass pots. EPN treated plants were obtained by adding 10 mL EPN solution (200 IJs/mL with 30mg EPN medium powder from Andermatt Biocontrol, CHE) to the bottom opening of the glass pots. Plants without EPNs were obtained by adding 10 mL of 3 mg/mL of EPN medium solution.

#### Petri dish assays

BHT attractiveness to WCR larvae was assessed in petri dish dual choice experiments (petri dish: 9 cm diameter, Greiner Bio-One GmbH, Frickenhausen, DE). A 5 mm layer of 1 % agarose (m/v, Sigma Aldrich Chemie, CHE) was poured into the dishes. Two to three 5 cm root pieces were placed at the two opposite sides of the petri dish. Two filter paper (90 mm, Whatman ^TM^, Sigma Aldrich Chemie, CHE) slices (length= 7 cm, width= 1 cm) were placed in parallel in between the root sections at four cm distance from each other. Five flash frozen healthy or EPN-infected WCR larvae (five days post-infection) were placed onto the filter paper slice respectively. BHT complementation was realized by adding 20 ng BHT in 100 μL 0.01% ethanol (≥99.8%, Sigma Aldrich Chemie, CHE) onto the filter paper slice. Control slices were imbibed with 100 μL 0.01% ethanol. Five third instar WCR larvae were given the choice between (i) control and BHT slices (n=20), (ii) healthy and EPN-infected larvae (n=10), (iii) healthy larvae and healthy larvae complemented with BHT (n=15) and (iv) EPN-infected larvae and healthy larvae complemented with BHT (n=15). The choice of WCR larvae was measured by adding five larvae in the center of the dish and recording their positions after 0.5 hr, 1.5 hr, 3 hr and 5 hr. WCR behavior towards EPN-infected WCR cadavers (frequency and duration of contact) was recorded over to hours using the same design (n=6). The preference of the six other insect species including *D. balteata* (n=19), *T. molitor* (n=16), *D. melanogaster* (n=14), *S. littoralis* (n=18), *P. xylostella* (n=16) and *G. mellonella* (n=10) to healthy and EPN-infected conspecifics was assessed using the same set-up.

### Root consumption by WCR larvae

Root consumption by WCR larvae over time was assessed in belowground olfactometers as described above. Root tissue were collected at 0, 24, 48, 72 and 96 hr (n_plant_=5, n_plant+WCR+EPN_=8) after adding WCR and EPNs. The difference between the root masses of plant+WCR+EPN complexes and healthy plants was used as a proxy for tissue removal. All collected roots were flash frozen for (*E*)-β-caryophyllene analyses (see below).

### EPN preference

BHT attraction to EPNs was evaluated in petri dish choice assays as described elsewhere (20). Briefly, a five mm layer of 0.5% agarose (Sigma Aldrich Chemie, CHE) was poured into the petri dishes. To test EPN behavioral response to BHT, control and BHT complemented exudates of *D. balteata* larvae were placed in two 5 mm diameter wells along the plate diameter and at 5 cm distance from each other. Exudates from *D. balteata* larvae were obtained by rinsing third instar larvae with tap water (50 μL per larva). BHT exudate complementation was performed by adding 20 ng BHT in 0.01% ethanol per well. Control exudates were obtained by adding the equivalent volume of 0.01% ethanol to the wells. BHT complementation of WCR larvae was realized by adding 20 ng BHT in 50 μL 0.01% ethanol onto flash frozen healthy WCR larvae. Control larvae were obtained by covering flash frozen healthy WCR larvae with 50 μL 0.01% ethanol. A third 5 mm diameter well was made in the center of the dishes to place sixty EPNs suspended in 100 μL water. EPNs were given a choice between: (i) control and BHT complemented insect exudates (n=20), (ii) healthy and EPN-infected larvae (n=20), (iii) healthy larvae and healthy larvae complemented with BHT (n=20) and (iv) EPN-infected larvae and healthy larvae complemented with BHT (n=20). The number of EPNs in each side was assessed 24 hr later.

### EPN infectivity and fecundity ability

EPN infection rate in belowground olfactometers was performed by collecting the larvae at 0, 24, 48, 72 and 96 hr (n=8) after adding WCR larvae and EPNs. The infection status of the WCR larvae was assessed visually.

The impact of BHT on EPN infectivity was evaluated in three experiments. First, the effect of BHT exposure on the resistance of WCR towards EPNs was tested by adding 50 μL of 0.4 ng/μL BHT in 0.01% ethanol (n=10) or 50 μL 0.01% ethanol only (n=10) on a slice of filter paper in a solo cup containing five third instar WCR. One day later, all the larvae were washed with 100% ethanol and tap water. Washed larvae were placed in new solo cups and 200 EPNs in 500 μL tap water were added. The resulting infection rate was recorded five days later.

Second, the effect of BHT exposure on EPN infectivity was tested by incubating EPNs in 0.2 ng/μL 0. 01% ethanol for 24 hr. After incubation, EPNs were washed twice with ethanol and tap water and then 500 EPNs were added into solo cups containing five third-instar WCR larvae (n=15). EPNs incubated in 0.01% ethanol were used as controls (n=15). The infection rate was recorded 5 days later.

Third, the impact of BHT release by EPN-infected cadavers on EPN predation success was tested in soil arenas. Six maize plants were sown in rectangular plastic trays (25 cm *11.5 cm * 9.5 cm, Migros, Bern, CHE), such as two sets of each three plants grew at about 15 cm distance from each other. After 12 days, 1500 EPNs in 2 mL tap water 0.01% ethanol containing 40 ng of BHT were added on one side, while 1500 EPNs in 2 mL tap water 0.01% ethanol only were added on the other side (n=12). Eight WCR larvae were placed in the middle section for four days. After this period, all larvae were collected, and the infection rate was recorded. The EPN-infected WCR larvae were collected to assess the number of emerging EPNs per larva as a proxy for EPN fecundity. Individual larvae were placed in adapted white traps (47). Briefly, the white trap consisted of 1.5 mL Eppendorf lid (Sarstedt AG & Co., Germany) placed upside down in a solo cup. The lid was covered with 2.5 cm diameter filter paper. Tap water was placed around the lid. Each larva was placed onto the filter paper and emerging EPNs could reach the water. The number of freshly emerged EPNs was counted 15 days after the emergence of the first EPNs.

### Volatile analysis

Plant (*E*)-β-caryophyllene emissions were measured using solid-phase micro-extraction-gas chromatography-mass spectrometry (SPME-GC-MS). Briefly, root tissues were ground in liquid nitrogen to a fine power and 100 mg was placed into 20 mL glass vials (Gerstel, Germany) for analysis as described below. Insect volatile profiles were also determined by GC-MS. All larvae were flash frozen prior to sampling and volatile profiles were obtained by placing five larvae of each treatment into a 20 mL glass vial. Gas chromatography analyses were performed using an Agilent 7820A GC interfaced with an Agilent 5977E MSD following protocols established by Erb *et al* (48) with a few modifications. Specifically, the SPME fiber (100 μm polydimethylsiloxane coating, Supelco, USA) was inserted into the vial for 30 min. The fiber was desorbed at 220 °C for 2 min. The column temperature was initially set at 60 °C for 1 min and increase to 200 °C at a speed of 5 °C min^−1^. The resulting GC-MS chromatograms were processed with Progenesis QI (informatics package from Waters, MA, USA) using default settings for spectral alignment and peak picking. Compound identification was realized using the NIST search 2.2 Mass Spectral Library and pure compound standards. Butylated hydroxytoluene (BHT) quantification was made using a standard curve of the pure compound (Sigma Aldrich Chemie, CHE) in 0.01% ethanol (v/v). (*E*)-β-caryophyllene (EBC) analysis was also modified from previous study (24). The protocol was similar as above described but the column temperature finally increased to 250 °C at a speed of 5 °C min^−1^.

### Data analysis

Preference data were analyzed by comparing the average difference between the proportion of WCR larvae or EPNs choosing control and treated sides to the null hypothesis H_0_=0 using analysis of variance (ANOVA). All other experiments were analyzed by ANOVAs followed by pairwise or multiple comparisons of Least Squares Means (LSMeans) and FDR-corrected post hoc tests (50). All analyses were carried out using R 3.2.2 (R Foundation for Statistical Computing, Vienna, Austria).

## ACKNOWLEDGEMENTS

We thank Marc Pfander for helping with data analysis, David Ermacora for rearing EPNs, Sarah Ettlin for helping with experiment, Matthias Erb, the Institute of Plant Sciences and the University of Bern for providing research infrastructure, and the gardeners of the Institute of Plant Sciences for their support with growing maize plants.

## AUTHOR CONTRIBUTIONS

C.A.M.R. designed the project. C.A.M.R. and X.Z. designed experiments. X.Z., R.A.R.M, C. vD., C.C.M.A., L.H. and C.A.M.R. carried out experiments. X.Z and C.A.M.R analyzed data. C.A.M.R. and X.Z. wrote the first draft of the manuscript. All authors contributed to the last version of the manuscript.

---

**Figure S1.**
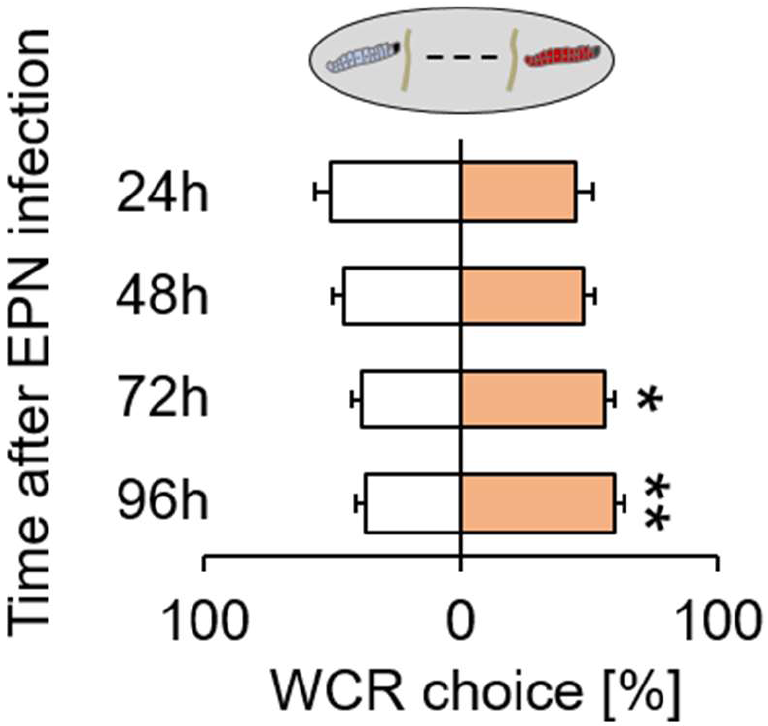
Nematode-infected cadavers become attractive at late infection stages. Proportions (mean ± SEM) of western corn rootworm (WCR) larvae choosing between uninfected cadavers and cadavers infected with entomopathogenic nematodes (EPNs) in belowground olfactometers. WCR choice was measured 24 hr, 48 hr, 72 hr, and 96 hr after infection (n=10-15). Stars indicate significant differences (*: p<0.05; **: p<0.01).

**Figure S2.**
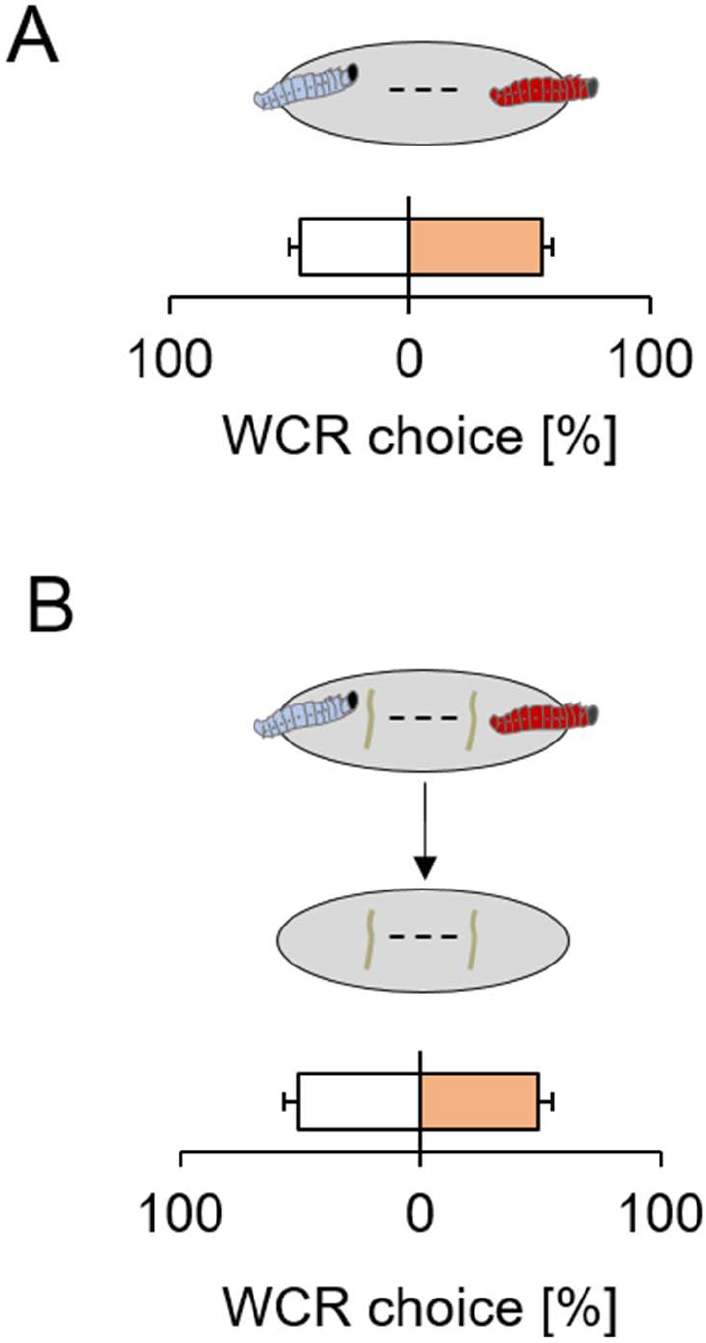
Attraction of the western corn rootworm to nematode-infected cadavers requires plant background odors. **A.** Proportions (mean ± SEM) of WCR choosing between uninfected cadavers and EPN-infected cadavers in petri dish assays without maize roots (n=20). **B.** Proportions (mean ± SEM) of WCR choosing between maize roots previously exposed to uninfected or EPN-infected cadavers in petri dish assays (n=20).

**Figure S3.**
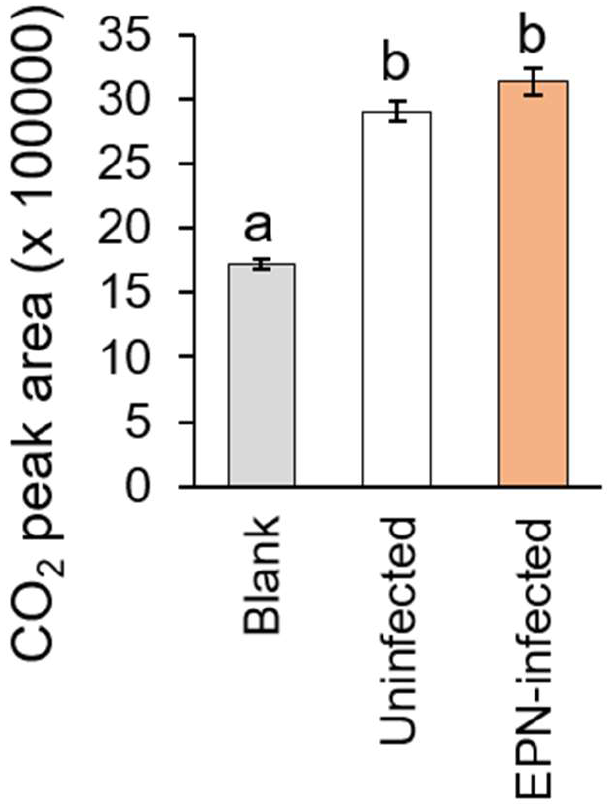
Infection by entomopathogenic nematodes does not alter CO_2_ emissions from western corn rootworm cadavers. CO_2_ content (mean peak area ± SEM) in an empty vial (Blank, n=8), in a vial containing five uninfected western corn rootworm (WCR) cadavers (n=12) or five WCR cadavers infected by entomopathogenic nematodes (EPNs; n=12). CO_2_ emissions were recorded by GC-FID over 20 min. Different letters indicate significant differences between treatments.

**Figure S4.**
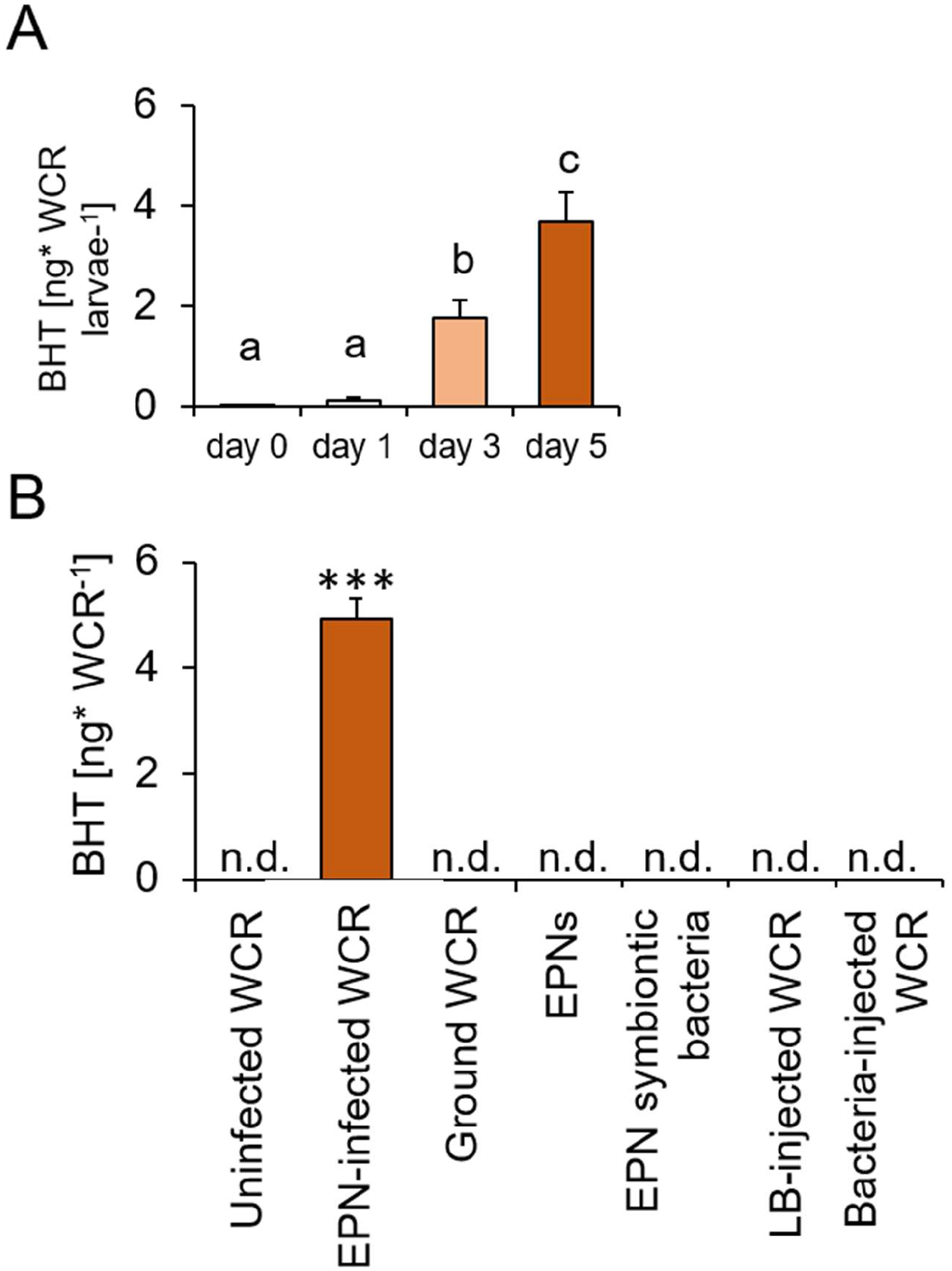
Butylated hydroxytoluene emission is specific to western corn rootworm infection by entomopathogenic nematodes. **A.** Butylated hydroxytoluene (BHT) emissions (mean ± SEM) of western corn rootworm (WCR) cadavers infected by entomopathogenic nematodes (EPNs) at 0, 1, 3 and 5 days after infection (n=4-5). Different letters indicate significant differences between treatments. **B.** BHT emissions (mean ± SEM) of uninfected WCR cadavers (n=8), EPN-infected WCR cadavers (n=5), ground uninfected WCR (n=3), EPNs (n=3), EPN symbiontic bacteria (n=3), bacteria-injected WCR (n=5) and bacterial medium (LB)-injected WCR (n=4). n.d.: not detected. Stars indicate significant differences (***: p<0.001).

**Figure S5.**
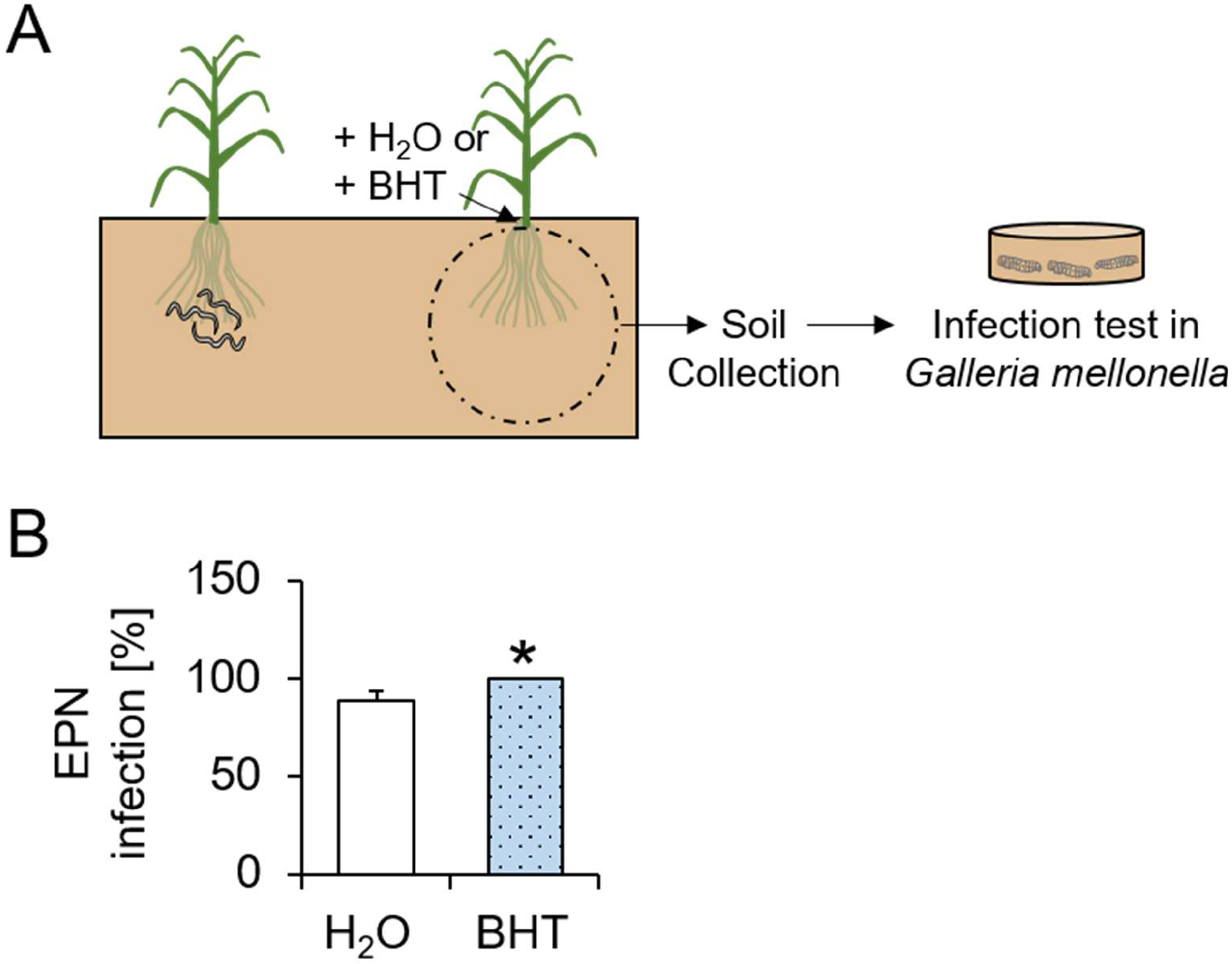
Butylated hydroxytoluene attracts entomopathogenic nematodes and increases their predation success in the soil. A. Visual representation of experimental setup. Each arena contained two pairs of plants separated by 15 cm. BHT or water (H_2_O) was added to the soil of one of the plant pairs. All treatment solutions contained 0.01% ethanol. EPNs were added to the soil of the second plant pair. After two days, 150 mg soil was collected from the BHT and H_2_O treated plants and placed in cups containing three *Galleria mellonella* larvae for infection tests. **B.** Proportion of EPN-infected *G. mellonella* larvae (mean ± SEM) exposed to soil collected from water (H_2_O) and BHT sides of the different arenas (n=12). Star indicate significant differences (*: p<0.05).

